# Reduced Hippocampal Inhibition and Enhanced Autism-Epilepsy Comorbidity in Mice Lacking Neuropilin 2

**DOI:** 10.1101/2021.06.11.448071

**Authors:** Carol Eisenberg, Deepak Subramanian, Milad Afrasiabi, Patryk Ziobro, Jack DeLucia, Pamela R. Hirschberg, Michael W. Shiflett, Vijayalakshmi Santhakumar, Tracy S. Tran

## Abstract

The neuropilin receptors and their secreted semaphorin ligands play key roles in brain circuit development by regulating numerous crucial neuronal processes, including the maturation of synapses and migration of GABAergic interneurons. Consistent with its developmental roles, the *neuropilin 2* (*Nrp2*) locus contains polymorphisms in patients with autism spectrum disorder (ASD). *Nrp2* deficient mice show autism-like behavioral deficits and propensity to develop seizures. In order to determine the pathophysiology in *Nrp2* deficiency, we examined the hippocampal numbers of interneuron subtypes and inhibitory regulation of hippocampal CA1 pyramidal neurons in mice lacking one or both copies of *Nrp2*. Immunostaining for interneuron subtypes revealed that *Nrp2*^*-/-*^ mice have reduced number of parvalbumin, somatostatin and Neuropeptide Y cells, mainly in CA1. Whole cell recordings identified reduced firing and hyperpolarized shift in resting membrane potential in CA1 pyramidal neurons from *Nrp2*^*+/-*^ and *Nrp2*^*-/-*^ mice compared to age-matched wild-type controls indicating decrease in intrinsic excitability. Simultaneously, the frequency and amplitude of spontaneous inhibitory postsynaptic currents (sIPSCs) are reduced in *Nrp2* deficient mice. A convulsive dose of kainic acid evoked electrographic and behavioral seizures with significantly shorter latency, longer duration and higher severity in *Nrp2*^*-/-*^ compared to *Nrp2*^*+/+*^ animals. Finally, *Nrp2*^*+/-*^ and *Nrp2*^*-/-*^, but not *Nrp2*^*+/+*^, mice have impaired cognitive flexibility demonstrated by reward-based reversal learning, a task associated with hippocampal circuit function. Together these data demonstrate a broad reduction in interneuron subtypes and compromised inhibition in CA1 of *Nrp2*^*-/-*^ mice, which could contribute to the heightened seizure susceptibility and behavioral deficits consistent with an ASD/epilepsy phenotype.

## Introduction

Epilepsy and autism spectrum disorders (ASD) exhibit a remarkable degree of comorbidity, suggesting shared pathological mechanisms ^1,2^. Developmental anomalies in inhibitory circuit formation can severely compromise the fine balance between neuronal excitation and inhibition (E/I) and have been proposed as candidate mechanisms linking childhood epilepsies and ASD ^3-6^. Nevertheless, the link between factors regulating formation of GABAergic circuits during development, their impact on cellular and synaptic inhibition and the risk for developing seizures and autism-like behavioral traits remains to be established.

Signaling of class 3 secreted semaphorins with their holoreceptors, Neuropilins (Nrp) and Plexins (Plxn) ^7,8^, are essential for wiring neural circuits by controlling neuronal morphogenesis, cell migration, axon guidance, synapse formation and survival ^9-13^. Nrp2 is expressed in inhibitory neuron precursor cells in the median ganglionic eminence (MGE) where it is regulated by the transcription factor Nkx2.1 and is essential for normal cortical interneuron migration ^14^. MGE-derived cortical/hippocampal neurons include parvalbumin (PV) and somatostatin (SOM) interneurons that mediate proximal and distal dendritic inhibition, as well as Neuropeptide Y (NPY) expressing neurons ^14-16^. PV neurons are numerically fewer and functionally impaired in developmental neuropsychiatric disorders including ASD and schizophrenia ^6,17^ while SOM and NPY neurons are reduced in epilepsy ^18^. Interestingly, mice with *Nrp2* deletion (*Nrp2*^*-/-*^) were reported to have reduced GABAergic neurons in hippocampal subfields ^19^ and were found to be impaired in social object recognition and social novelty preference ^20^ consistent with a potential link between reduction in interneurons and ASD-like phenotype. Importantly, mice with a single copy of *Nrp2* (*Nrp*^*+/-*^) show increased susceptibility to induction of behavioral seizures while mice with both copies deleted (*Nrp2*^*-/-*^) mice were reported to exhibit seizures on handling ^19^ suggesting that loss of Nrp2 may also contribute to developmental epilepsy. Although the association between Nrp2 and human epilepsy is unclear ^21^, Nrp2 polymorphism is associated with autism in two patient populations ^22,23^. Moreover, deletion of the Nrp2 ligand, semaphorin 3F (Sema3F), in GABAergic neurons resulted in increased seizure susceptibility ^24^ and development of autism-like behaviors ^25^, suggesting that mice with deficient Nrp2 signaling could serve as a model to assess inhibitory circuit dysregulation associated with the propensity for autism/seizure phenotypes.

The hippocampal formation is a major focus of histological and electrophysiological abnormalities in temporal lobe epilepsy and is functionally compromised in rodent models with asocial/autistic behaviors ^26-29^. Nrp2 plays a role in maturation of both glutamatergic and GABAergic circuits in the hippocampus. Nrp2 is expressed in hippocampal projection neurons where it regulates morphogenesis and excitatory synaptic function ^30-32^. Consistent with its role in interneuron migration, hippocampal GABAergic neurons, particularly NPY neurons, were found to be reduced in *Nrp2* deficient mice ^19^. Curiously, although PV and SOM neurons derive from the MGE and require Nrp2 for cortical migration ^14,19^, limited loss of PV neurons and no reduction in SOM neurons were found in hippocampal subfields of *Nrp2*^*-/-*^ mice. This contrasts with the strong deficit in hippocampus dependent behavioral tasks and propensity to seizures in *Nrp2* mutant mice ^19,20^. While significantly higher mEPSC frequencies, with no difference in amplitude, from hippocampal granule cells were reported in *Nrp2*^*-/-*^ as compared to *Nrp2*^*+/+*^ littermate controls ^30^, functional analysis of intrinsic physiology and inhibitory regulation of hippocampal principal neurons are needed to better inform the underlying circuit changes in these animals. Additionally, while *Nrp2*^*+/-*^ mice failed to exhibit the significant behavioral deficits observed in *Nrp2*^*-/-*^ mice ^20^, both *Nrp2*^*+/-*^ and *Nrp2*^*-/-*^ mice were reported to have comparable reduction in GABAergic neurons ^19^, suggesting that the reduction in interneurons and behavioral outcomes may be differentially impacted by the extent of Nrp2 deletion. However, whether *Nrp2*^*+/-*^ and *Nrp2*^*-/-*^mice show differences in cellular and synaptic inhibition and seizure susceptibility has not been examined. This study was conducted to determine the effect of heterozygous and homozygous *Nrp2* deletion on intrinsic physiology and inhibitory inputs to hippocampal CA1 pyramidal cells and assess whether there are genotype-dependent increases in seizure susceptibility and memory/learning deficits associated with ASD phenotypes.

## Materials and Methods

### Animals

All experiments were performed in accordance with IACUC protocols approved by Rutgers University, Newark, NJ, and the University of California at Riverside, CA and in keeping with the ARRIVE guidelines. *Neuropilin 2* (*Nrp2*) knockout mouse line has been described previously in detail ^33^. Mice used in this study were backcrossed for at least 10 generations to the C57BL/6NTac background strain. *Nrp2* genotypes were confirmed by using polymerase chain reaction (PCR).

### Immunostaining

*Nrp2*^*+/+*^ (*wild-type*), *Nrp2*^*+/-*^ (*heterozygote*) and *Nrp2*^*-/-*^ (*homozygote*) mice (3-5 months) were subjected to transcardial perfusion with 4% paraformaldehyde (PFA). Brains were dissected and postfixed for 2 hours fixation with 4% PFA then imbedded in OCT medium. Brains were sectioned on a cryostat at 20 μm thickness at 200 μm intervals through the anterior to posterior extent of the hippocampus and cortex along the coronal plane. Sections were processed for immunocytochemistry as previously described ^34^, with the following modification. Coverslips were mounted on microscope slides using Mowiol (cat. No. 81381 Aldrich), plus 10% p-Phenylenediamine (PPD, cat No. 78460) anti-fade mounting media. Primary antibodies used were: mouse monoclonal anti-Parvalbumin (1.5:500, Swant, PV235); mouse monoclonal anti-Parvalbumin (1:750, Sigma, P3088); rabbit polyclonal anti-Neuropeptide Y (1:1000, Abcam, ab30914); rat monoclonal anti-Somatostatin (1:150, Millipore, MAB354) and for visualization, AlexaFluor 488 or Cy5 (1:500, Jackson Immuno Research Laboratories); AlexaFluor 546 (1:500 Invitrogen).

### Cell Counts and Data Analysis

For cells containing parvalbumin (PV), Neuropeptide Y (NPY) and somatostatin (SOM), the soma was counted in 4 major regions: Dentate gyrus/hilar region/CA4 (DG) region, CA1, CA2 and CA3 regions (see Supplementary Fig. 1 for demarcation) modified from ^19^. Within these demarcated regions, PV, NPY and SOM-stained cells were quantified in the entire hemi-hippocampus. Ten PV, NPY, SOM and Dapi stained hemi-hippocampal sections (at 200 µm intervals) per animal were counted using the ImageJ (Fiji) software. Experimenters, total of two whom independently counted the cells, were blinded to genotype. Average measures per animal were used for statistical analysis (n= at least 3 animals/genotype). The total number of PV, NPY and SOM cells were counted in the hippocampal region in each of 10 sections. Total counts were averaged together to establish a single value of cell counts per hemi-hippocampal region for each mouse.

### Photo-documentation

All immunocytochemically processed brain sections were imaged using the Zen software (Zeiss) and Zeiss 510 LSC Confocal System. Representative nonoverlapping fields were imaged and stitched together to reconstruct hippocampal and cortical sections. Compiled z-stack images were exported as tif files. Images were captured with EC Plan-Neofluar 10X/0.30 M27, and Plan-Neofluar 40x/1.3 Oil DIC and the following laser lines: 405, 488, 543, 633 nm.

### Slice Physiology

Horizontal hippocampal slices (350 µm) were prepared from male and female 4-8 weeks old *Nrp2*^*+/+*^, *Nrp2*^*+/-*^ and *Nrp2*^*-/-*^ mice following euthanasia under isoflurane anesthesia ^35,36^. Slices were prepared in ice-cold high Sucrose artificial Cerebro-Spinal Fluid (Sucrose-aCSF) that contained (in mM) 85 NaCl, 75 sucrose, 24 NaHCO_3_, 25 glucose, 4 MgCl_2_, 2.5 KCl, 1.25 NaH_2_PO_4_, and 0.5 CaCl_2_ using Leica VT1000 or VT1200S Vibratomes (Wetzlar, Germany). Sliced were bisected and incubated at 32 ± 1°C for a minimum of 1 hour in a submerged holding chamber containing recording aCSF (in mM) 126 NaCl, 2.5 KCl, 2 CaCl_2_, 2 MgCl_2_, 1.25 NaH_2_PO_4_, 26 NaHCO_3_ and 10 D-glucose and subsequently held at room temperature (RT). All solutions were saturated with 95% O_2_ and 5% CO_2_ and maintained at a pH of 7.4 for 2-6 h ^35,36^.

Slices were transferred to a submerged recording chamber and perfused with oxygenated aCSF at 33 ± 1°C and CA1 pyramidal cells were patched under IR-DIC visualization using Nikon Eclipse FN-1 (Nikon Corporation, Shinagawa, Tokyo, Japan) or Olympus BX50 (Olympus Corporation, Shinjuku, Tokyo, Japan) microscope, using 40X water-immersion objectives and borosilicate microelectrodes (3–5 MΩ). Whole-cell voltage- and current-clamp recordings were obtained using Axon Instruments Axopatch 200B or MultiClamp 700B amplifiers (Molecular Devices, Sunnyvale, CA). Data were low-pass filtered at 3 kHz, digitized using DigiData 1440A and acquired using pClamp10 at 10-kHz sampling frequency. The internal solution for recordings contained (in mM) 125 KCl, 10 K-gluconate, 10 HEPES, 2 MgCl2, 0.2 EGTA, 2 Na-ATP, 0.5 Na-GTP, 10 PO Creatine and 0.3% Biocytin and was titrated to a pH 7.25 with KOH (270–290 mOsm). Recorded neurons were held in current clamp at -60 mV to elicit voltage responses to 500 milli-second positive and negative current steps. Active and passive properties including spike frequency, action potential amplitude, spike frequency adaptation ratio, resting membrane potential, input resistance and sag ratio were measured and calculated as described previously ^35,37^. Neurons were then held in voltage clamp at -60 mV to record sIPSCs which were isolated by inclusion of the glutamate receptor antagonist, kynurenic acid (3 mM KyA, Tocris, Ellisville, MO) in the recording aCSF. Individual sIPSCs were detected using custom software in IgorPro7.0 (WaveMetrics’, Lake Oswego’, OR) and different parameters were analyzed. Events were visualized’, and any “noise” that spuriously met trigger specifications was rejected. Cumulative probability plots of sIPSC parameters were constructed using IgorPro by pooling an equal number of sIPSCs from each cell ^36^.

### In vivo video-EEG recordings

Seventeen mice (6 *Nrp2*^*+/+*^, 6 *Nrp2*^*+/-*^ and 5 *Nrp2*^*-/-*^) were surgically implanted with a tungsten wire electrodes (50 µm, California Fine Wire company), placed over the granule cell layer of DG (AP:2 mm, ML: 1.2-1.5 mm, DV: 1.7 mm from bregma; see fig X) guided by periodic electrical stimulation of the angular bundle (AP: 0.5 mm, ML:3.2 mm, DV: 0.8-1 mm). Two additional screw electrodes (invivo1) on the contralateral side served as ground and reference. After 3-5 days of recovery, mice were connected to a tethered video EEG monitoring system and baseline recordings were obtained for 30 minutes. Signals were sampled at 10kHz, amplified (x100, 8202-SE3, Pinnacle Technologies), digitized (Powerlabs16/35, AD Instruments) and recorded using LabChart 8.0 (AD instruments). Following baseline recordings, mice received a single high dose of KA (25mg/kg) and their latency to electrographic seizure, seizure duration and severity were measured. Seizures were defined as rhythmic activity exceeding a threshold of mean baseline amplitude +2.5 standard deviation for >5 seconds ^38^. Artifacts due to electrical, exploratory and grooming behavior were identified and removed manually from the raw EEG’s before analysis. Seizures were scored based on a modified Racine scale ^39^ as: Stage-1, absence-like immobility; Stage-2, hunching with facial or manual automatisms; Stage-3, rearing with facial or manual automatisms and forelimb clonus; Stage-4, repeated rearing with continuous forelimb clonus and falling; and Stage-5, generalized tonic clonic convulsions with lateral recumbence or jumping and wild running followed by generalized convulsions ^40^. Seizure severity scores in the first 30 minutes were averaged over 10 min epochs. A cumulative seizure score was used to quantify overall seizure severity.

### Reversal learning behavioral procedure

A total of 16, 3-5 months old mice were used for behavioral testing: 6 *Nrp2*^*-/-*^, 5 *Nrp2*^*+/-*^ and 5 *Nrp2*^*+/+*^. Mice were placed on a restricted food diet of approximately 2 g of standard chow each day. The chow (Purina, St. Louis MO, USA) was given in their home cage after behavioral procedures were complete. Animals were weighed daily, and their body weights were maintained to 85-90% of their original weight. Mice were tested in eight operant conditioning chambers (Med Associates, St. Albans VT, USA) as previously described ^34^. Operant chamber operation and data collection were carried out with Med Associates proprietary software (Med-PC V). While placed on food restriction, mice were simultaneously habituated to the operant conditioning chamber for one 15-minute session. The next day, mice were conditioned to find pellets in the food cup in a 20-minute session in which food pellets were dispensed on a random-time 60 sec schedule. Levers were retracted during this phase. The next day mice were trained to use levers. During each session a single lever was inserted into the operant conditioning chamber. Each lever press resulted in delivery of a single 20-mg grain-based chocolate flavored food pellet (Bio-serv, Frenchtown NJ USA) into the food cup. The session terminated after either 20 lever presses or 20 minutes elapsed from the start of the session. The mice completed two training sessions per day, one session with the left lever and one session with the right lever.

After all mice demonstrated proficiency in instrumental conditioning, they were placed on a reversal-learning task. In this phase both levers were extended simultaneously. One lever was designated as the rewarded lever, and the other lever as the non-rewarded lever. For each trial, responses on the rewarded lever always produced a food pellet, and responses on the non-rewarded lever never delivered a pellet. When no pellet was attained, both levers were retracted for 3 sec and reinserted. After the mouse earned between 10 and 14 food pellets, the contingencies assigned to each lever were reversed; responses on the previously rewarded lever produced no pellets, whereas responses on the non-rewarded lever produced pellets. Mice remained in the operant chamber until 80 trials were completed/day. Task performance was evaluated by win/stay and lose/shift ratios for each animal. The win/stay ratio is calculated as win-stay divided by the total of win-stay and win-shift choices. Similarly, the lose/shift ratio is calculated as lose-shift divided by the total of lose-stay and lose-shift choices.

### Statistical Analysis

One-way or repeated-measure ANOVA (SigmaPlot) was used to test for statistical differences between the active and passive neuronal properties and IPSC parameters in electrophysiological recordings. For slice physiological recordings, the sample size is the number of cells and the number of mice they were obtained from is reported as well. The two-way ANOVA and post hoc Tukey’s multiple comparison correction (Graph Pad Prism Software) were used for statistical differences in the cell counts and reversal-learning task. Data that failed normality was analyzed using appropriate nonparametric tests. Significance was set to *p*<0.05. Data are shown as mean ±SEM or median and interquartile range as appropriate.

## Results

### Developmental Loss of Nrp2 significantly decreases the number of specific interneuron populations in the hippocampus

Nrp2 has been proposed to contribute to cortical and striatal interneuron numbers by regulating the migration patterns of MGE-derived interneuron progenitors ^14,41^. However, previous studies have reported a limited reduction in interneuronal numbers, restricted to NPY+ interneuronal subtype, in the hippocampal CA1 region of *Nrp2*^*-/-*^ mice ^19^. Therefore, we first asked whether the number of MGE-derived interneuron subtypes in the hippocampal CA1 is altered in *Nrp2*^*+/-*^ and *Nrp2*^*-/-*^ compared to *Nrp2*^*+/+*^ littermates. We counted the numbers of specific interneuron populations using set parameters taken from ten coronal brain sections (at 20 μm thickness) obtained at 200 μm intervals through the entire anterior-posterior hemi-hippocampus (Fig. S1 and see Methods).

First, we focused on the PV+ interneurons, which include soma-targeting fast-spiking basket cells, axo-axonic cells targeting the axon initial segment of principal cells, as well as classes of interneurons projecting to pyramidal cell dendrites ^42,43^. We found significant decreases in PV+ cell populations (Fig. 1A-E) between *Nrp2*^*-/-*^ versus *Nrp2*^*+/+*^, 38.2% in CA1 and 39.3% in CA2 (Fig. 1F; cell numbers in CA1: 24.18±2.33 *Nrp2*^*+/+*^, 21.62±1.35 *Nrp2*^*+/-*^, 14.95±1.14 *Nrp2*^*-/-*^; *F*_(2,24)_=11.13 two-way ANOVA with Tukey’s multiple comparison correction *p<*0.0001; cell numbers in CA2: 10.65±0.06 *Nrp2*^*+/+*^, 10.44±1.34 *Nrp2*^*+/-*^, 6.46±1.36 *Nrp2*^*+/+*^; F_(2,24)_=11.13 two-way ANOVA with Tukey’s multiple comparison correction,*p*=0.0435) as compared to previous report of a 27% reduction ^19^. The overall reduction in PV+ cells (Fig. 1G; total mean cell number: 48.44±4.8 *Nrp2*^*+/+*^, 47.55±2.76 *Nrp2*^*+/-*^, 36.06±2.50 *Nrp2*^*-/-*^; *F*_(2,24)_=11.13 two-way ANOVA with Tukey’s multiple comparison correction *p*=0.0004) was 25.6% in all hippocampal regions in *Nrp2*^*-/-*^ as compared to *Nrp2*^*+/+*^, with no difference between *Nrp2*^*+/+*^ versus *Nrp2*^*+/-*^ animals.

Next we examined NPY+ cells, which were previously reported to be decreased in the *Nrp2*^*-/-*^ CA1 as compared to *Nrp2*^*+/+*^ littermates ^19^. Our systematic analysis identified a 28.5% reduction in NPY+ neurons selectively in the CA1 region of *Nrp2*^*-/-*^ mice as compared to *Nrp2*^*+/+*^ littermates (Fig. 2A-E), with no statistically significant differences detected in other hippocampal regions (Fig. 2F; cell numbers in CA1: 38.80±1.73 *Nrp2*^*+/+*^, 35.27±3.83 *Nrp2*^*+/-*^, 27.75±0.75 *Nrp2*^*-/-*^; *F*_(2,24)_=8.017 two-way ANOVA with Tukey’s multiple comparison correction *p*=0.003). While there is an overall 24% reduction in NPY+ neurons of *Nrp2*^*-/-*^ versus *Nrp2*^*+/+*^ littermates when all hippocampal subregions were combined (Fig. 2G; total mean cell number: 89.43±5.81 *Nrp2*^*+/+*^, 87.75±5.74 *Nrp2*^*+/-*^, 67.98±3.46 *Nrp2*^*-/-*^; *F*_(2,24)_=8.017 two-way ANOVA with Tukey’s multiple comparison correction *p*=0.0040), differences between *Nrp2*^*+/-*^ versus *Nrp2*^*+/+*^ animals failed to reach statistical significance (Fig. 2G).

**Figure 1.**
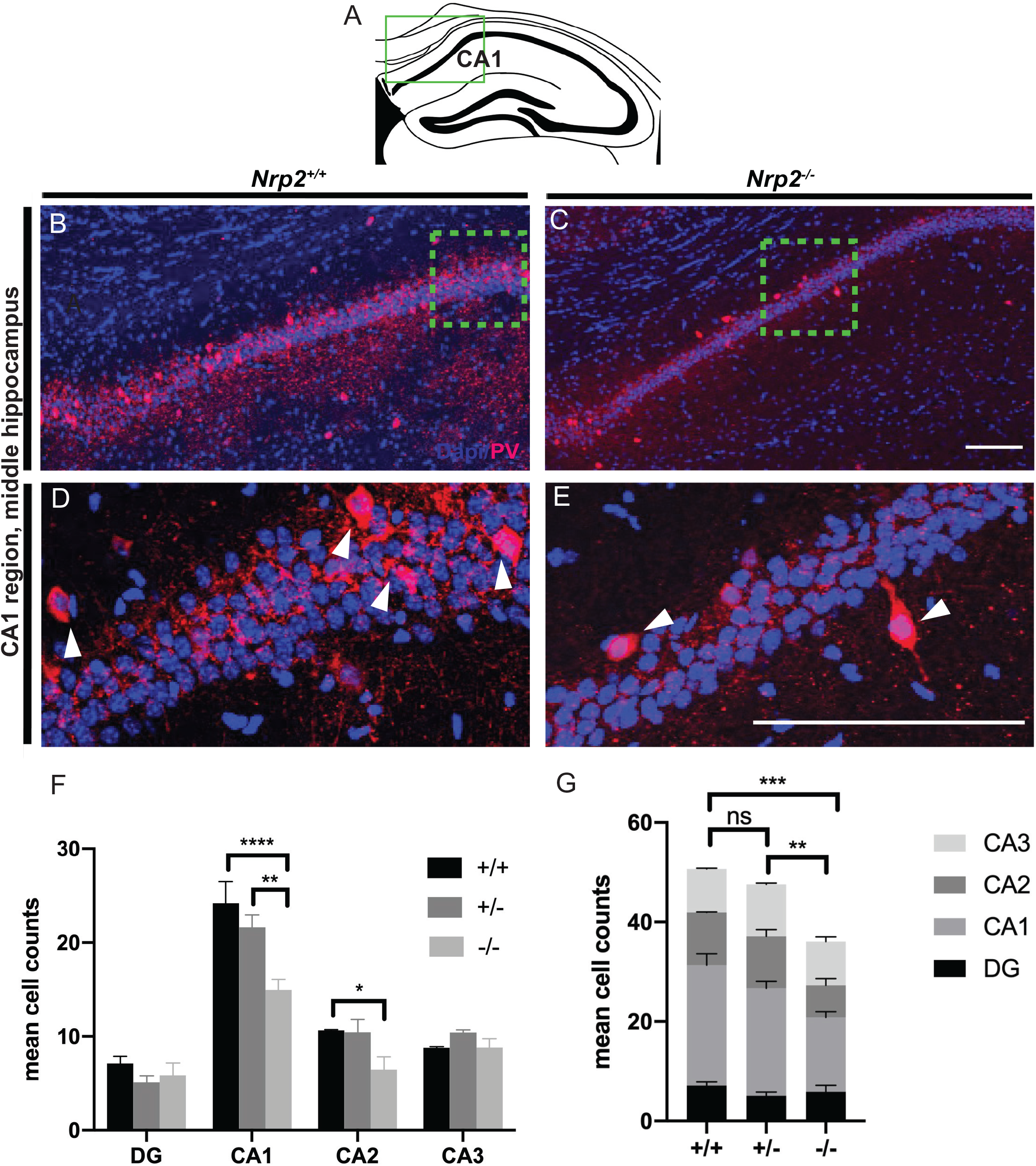
Developmental deletion of *Nrp2* leads to reduced numbers of parvalbumin (PV) expressing neurons in the CA1 region of the hippocampus. A) Schematic of a coronal adult hippocampal section, green box indicates CA1 hippocampal region of images in B-E. B-C) Immuno-labeled of *Nrp2*^*+/+*^ (littermate control) and *Nrp2*^*-/-*^ brain sections, respectively, with anti-PV (red) and Dapi (blue). D-E) High magnification images of green boxes in B and C, respectively. Scale bars: 100μm in C and E for B and D, respectively. F) Quantification of the averaged number of PV+ cells by hippocampal region. Significantly fewer number of PV+ neurons found in CA1 region of *Nrp2*^*-/-*^ compared to *Nrp2*^*+/+*^ mice. Error bars are ± SEM; two-way ANOVA, post-hoc Tukey: \*\*\*\**p<*0.0001 +/+ vs. -/-; \*\**p*=0.0012 +/- vs. -/- in CA1; \**p*=0.0435 +/+ vs. -/- in CA2. G) Quantification of total mean PV+ cells. Significantly more PV expressing neurons found in hippocampal region of *Nrp2*^*+/+*^ and *Nrp2*^*+/-*^ mice compared to *Nrp2*^*-/-*^ mice. Error bars are ± SEM; two-way ANOVA, post-hoc Tukey: \*\*\**p*=0.0004 +/+ vs. -/-; \*\**p*=0.0048 +/- vs. -/-; ns *p*=0.6072. For F and G: n=3 animals/genotype.

**Figure 2.**
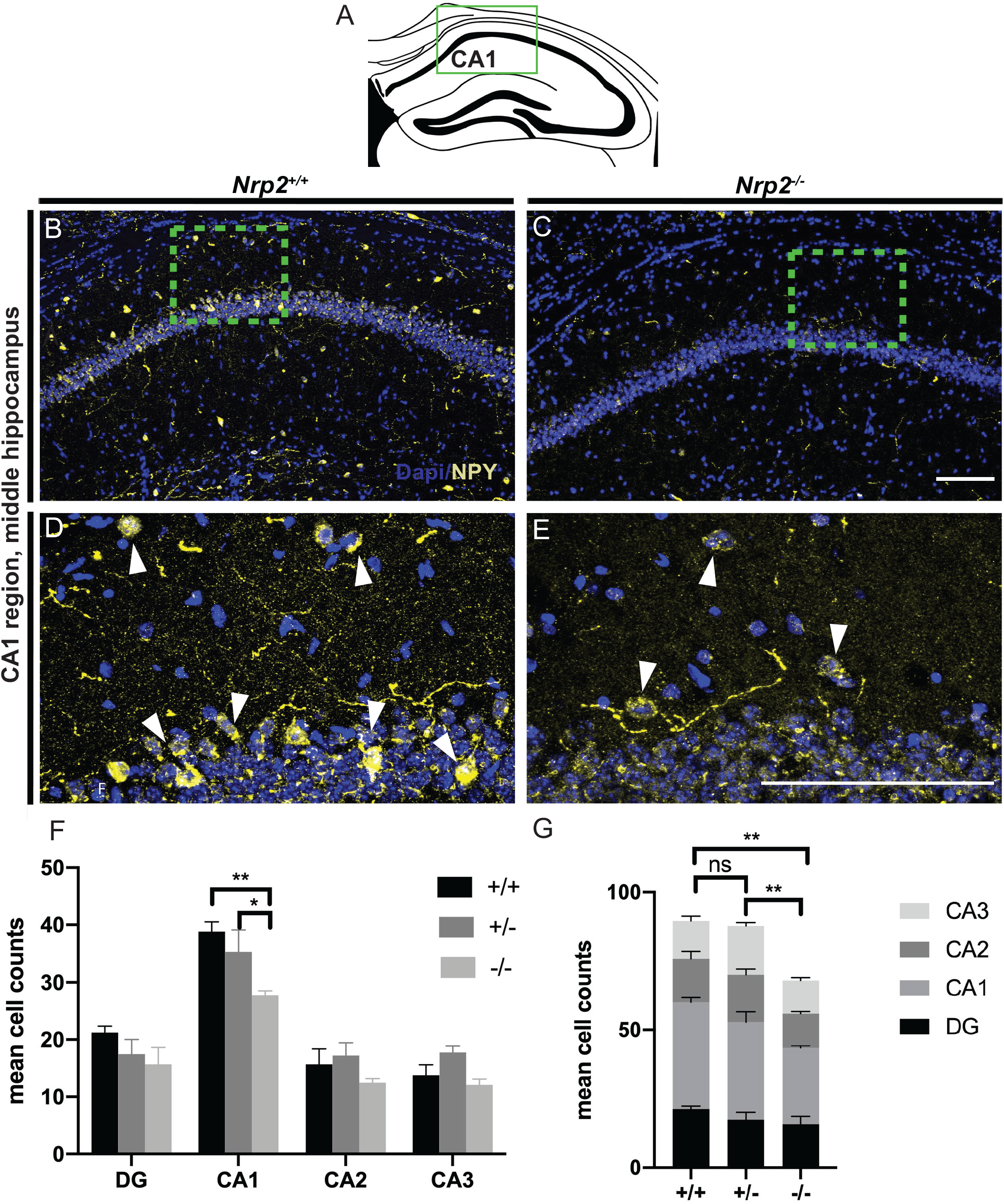
*Nrp2* deletion results in reduced numbers of neuropeptide Y (NPY) expressing neurons in the CA1 region. A) Schematic of a coronal adult hippocampal section, green box indicates CA1 hippocampal region of images in (B-E). B-C) Immuno-labeled *Nrp2*^*+/+*^ and *Nrp2*^*-/-*^ brain sections, respectively, with anti-NPY (yellow) and Dapi (blue). D and E) High magnification images of green boxes in B and C, respectively, with white arrows indicating cells. Scale bars: 100μm in C and E for B and D, respectively. F) Quantification of the averaged number of NPY+ cells by hippocampal region. Significantly more NPY+ neurons found in CA1 region of *Nrp2*^*+/+*^ compared to *Nrp2*^*-/-*^ mice. Error bars are ± SEM; two-way ANOVA, post-hoc Tukey: \**p*=0.0474, \*\**p*=0.0030. G) Quantification of total mean NPY+ cells. Significantly fewer NPY expressing neurons found in hippocampal region of *Nrp2*^*-/-*^ compared to *Nrp2*^*+/+*^ or *Nrp2*^*+/-*^ mice. Error bars are ± SEM; two-way ANOVA, post-hoc Tukey: \*\**p*=0.0040 +/+ vs. -/-, \*\**p*=0.0078 +/- vs. -/-, and ns *p*=0.9569. In F and G: n=3 animals/genotype.

Finally, we examined SOM+ neurons, which specifically target dendrites and facilitate their inhibition. In contrast with previous report of no change in SOM+ cell density ^19^, we found a striking reduction in SOM+ neurons in all hippocampal subfields of the *Nrp2*^*-/-*^ mutants as compared to littermate controls (Fig. 3). We observe a 37.1% and 42.7% decrease in SOM+ neurons in CA1 and CA2, respectively, from *Nrp2*^*-/-*^ as compared to *Nrp2*^*+/+*^ mice (Fig. 3A, B, E-L, U; cell numbers in CA1: 38.35±2.75 *Nrp2*^*+/+*^, 38.06±3.12 *Nrp2*^*+/-*^, 24.12±0.77 *Nrp2*^*-/-*^; *F*_(2,24)_=41.26 two-way ANOVA with Tukey’s multiple comparison correction *p*<0.0001; cell numbers in CA2: 17.71±0.24 *Nrp2*^*+/+*^, 13.06±1.69 *Nrp2*^*+/-*^, 10.15±0.85 *Nrp2*^*-/-*^; *F*_(2,24)_=41.26 two-way ANOVA with Tukey’s multiple comparison correction *p*=0.0193), with greater loss of SOM+ neurons toward the posterior hippocampus. We see an even larger reduction of SOM+ neuron in both the CA3 (Fig. 3C, M-P) and dentate gyrus (Fig. 3D, Q-T) at a 61.2% and 54.2% loss in *Nrp2*^*-/-*^ as compared to *Nrp2*^*+/+*^ mice, respectively (Fig. 3U; cell numbers in CA3: 23.55±1.20 *Nrp2*^*+/+*,^ 17.29±2.82 *Nrp2*^*+/-*^, 9.13±0.59 *Nrp2*^*-/-*^; *F*_(2,24)_=41.26 two-way ANOVA with Tukey’s multiple comparison correction *p*=0.0001; cell numbers in DG: 18.65±1.54 *Nrp2*^*+/+*^, 12.90±2.28 *Nrp2*^*+/-*^, 8.53±1.04 *Nrp2*^*-/-*^; *F*_(2,24)_=41.26 two-way ANOVA with Tukey’s multiple comparison correction *p*=0.0018). The overall decrease in SOM+ neurons (Fig. 3V; total mean cell number: 98.26±2.06 *Nrp2*^*+/+*^, 81.31±9.29 *Nrp2*^*+/-*^, 51.93±2.39 *Nrp2*^*-/-*^; *F*_(2,24)_=41.26 two-way ANOVA with Tukey’s multiple comparison correction *p*<0.0001 for *Nrp2*^*-/-*^ vs. *Nrp2*^*+/+*^, *p*=0.0085 for *Nrp2*^*+/-*^ vs. *Nrp2*^*+/+*^) was 47.2% in all hippocampal regions in *Nrp2*^*-/-*^ as compared to *Nrp2*^*+/+*^, with a smaller but significant reduction of 17.2% between *Nrp2*^*+/-*^ versus *Nrp2*^*+/+*^ animals. Together, we find that all three major interneurons populations, PV+, NPY+ and SOM+, are significantly reduced in the CA1 of *Nrp2*^*-/-*^ mice as compared to *Nrp2*^*+/+*^ controls. This finding is in contrast to a previous report showing the majority of the interneuron loss in the NPY+ population and a slight reduction in PV+ neurons in CA1 and CA3 with no loss of SOM+ neurons ^19^. Furthermore, we demonstrate that SOM+ neurons show the greatest decrease in all hippocampal subfields.

**Figure 3.**
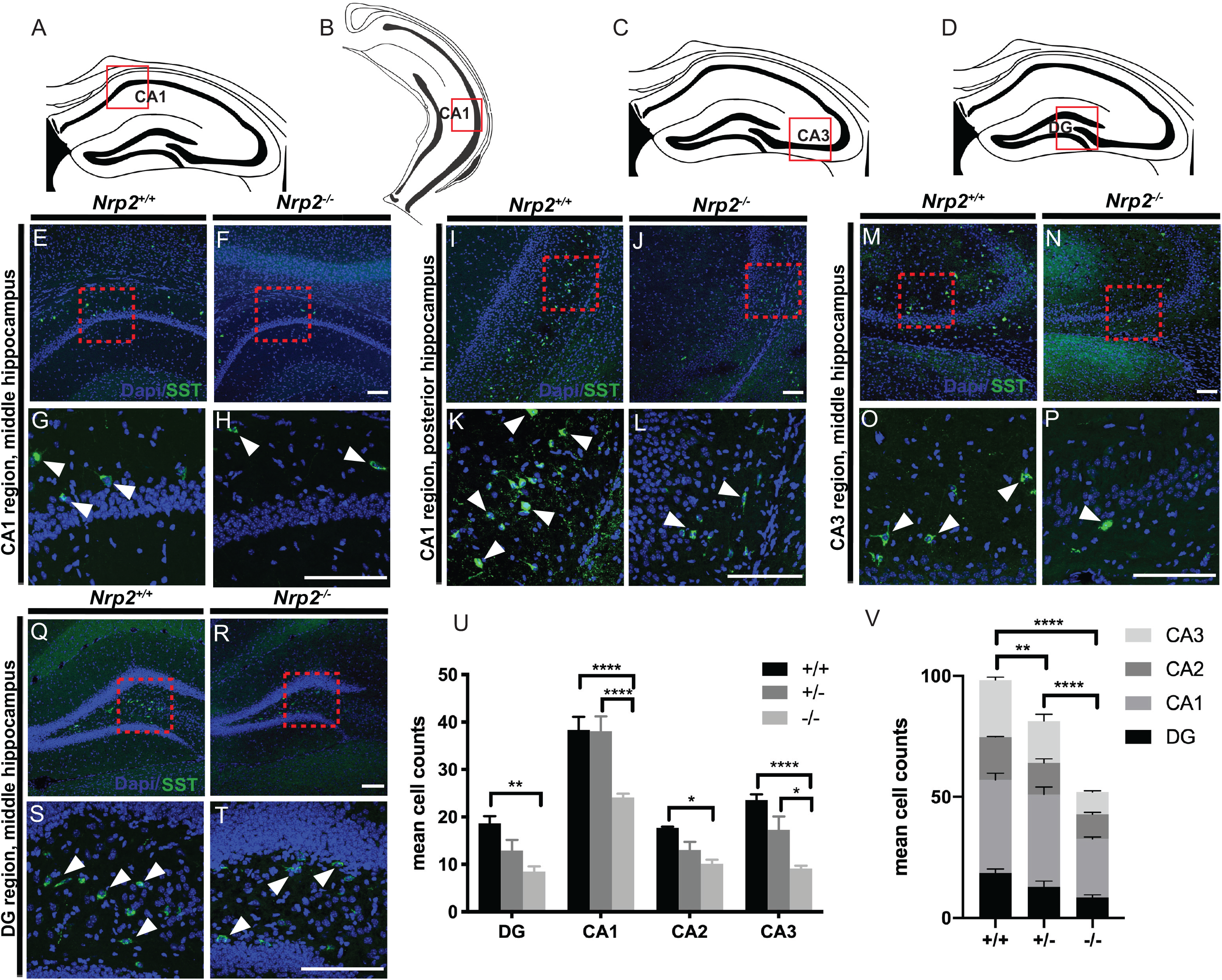
Somatostatin (SOM) expressing interneurons in CA1, CA3 and dentate gyrus (DG) hippocampal regions are significantly fewer in number in *Nrp2*^*-/-*^ compared wild type (*Nrp2*^*+/+*^). A-B) Schematics show coronal adult sections of the CA1 region in the middle and posterior hippocampus, respectively. Red box in A and B indicates the CA1 region illustrated by images in E-H and I-L, respectively. C-D) Schematics of coronal adult sections in the middle hippocampus, red boxes indicate the CA3 and DG region of images in M-P and Q-T, respectively. E-H) Immuno-labeled of littermates *Nrp2*^*+/+*^ (E, G) and *Nrp2*^*-/-*^ (F, H) sections with anti-SOM (green) and Dapi (blue) in the middle CA1 region. Red boxes in E and F are the locations of the high magnification images shown in G and H, respectively. I-L) Immuno-labeled of littermates *Nrp2*^*+/+*^ (I, K) and *Nrp2*^*-/-*^ (J, L) sections with anti-SOM (green) and Dapi (blue) in the posterior CA1 region. Red boxes in I and J are the locations of the high magnificent images shown in K and L, respectively. M-P) Immuno-labeled sections from littermates *Nrp2*^*+/+*^ (M, O) and *Nrp2*^*-/-*^ (N, P) with anti-SOM (green) and Dapi (blue) in the CA3 region. Red boxes in M and N show the location of high magnification images in O and P, respectively. Q-T) Littermates *Nrp2*^*+/+*^ (Q, S) and *Nrp2*^*-/-*^ (R, T) sections are immuno-labeled with anti-SOM (green) and Dapi (blue) in the DG region. Red boxes in Q and R show the location of high magnification images in S and T, respectively. Scale bars: 100 μm in F, J, N and R for E-F, I-J, M-N and Q-R, respectively; 100 μm in H, L, P and T for G-H, K-L, O-P and S-T, respectively. U) Quantification of the averaged number of SOM+ cells by hippocampal region. Significantly fewer number of SOM+ neurons found in CA1, CA3 and DG regions of *Nrp2*^*-/-*^ compared to *Nrp2*^*+/+*^ mice. Error bars are ± SEM; two-way ANOVA, post-hoc Tukey: \*\**p*=0.0018 +/+ vs. -/- in DG; *\*\*\*\*p*<0.0001 +/+ vs. -/- and +/- vs. -/- in CA1; \**p*=0.0193 +/+ vs. -/- in CA2; *\*\*\*\*p*<0.0001 +/+ vs. -/-, \**p*=0.0113 +/- vs. -/- in CA3. V) Quantification of total mean SOM+ cells. Significantly fewer SOM+ expressing neurons overall found in the hippocampus of *Nrp2*^*+/-*^ and *Nrp2*^*-/-*^ as compared to *Nrp2*^*+/+*^ mice. Error bars are ± SEM; two-way ANOVA, post-hoc Tukey: \*\*\*\**p*<0.0001 +/+ vs. -/- and +/- vs. -/-; \*\**p*=0.0085 +/+ vs. +/-. For U and V, n=3 animals/genotype.

### Synaptic Inhibition is reduced in mice lacking Nrp2

In light of the marked decrease in all three interneuronal classes in hippocampal CA1 of *Nrp2*^*-/-*^ mice, we examined whether the deletion of Nrp2 alters synaptic inhibitory inputs to CA1 pyramidal cells (Fig. 4A-D). In voltage clamp recordings from CA1 pyramidal cells, the frequency of sIPSCs in Nrp2 deficient mice was significantly reduced compared to *Nrp2*^*+/+*^ littermates but was not different between *Nrp2*^*+/-*^ and *Nrp2*^*-/-*^ mice (Fig. 4C: frequency in Hz, *Nrp2*^*+/+*^: 23.91±2.30, median=24.41, IQR=6.37-30.12, n=12 cells/ 4 mice; *Nrp2*^*+/-*^: 16.45±2.44, median=8.83, IQR=3.378-20.33, n=13 cells/ 4 mice; *Nrp2*^*-/-*^: 17.57±1.32, median=8.753, IQR=4.089-30.192, n=19 cells/ 5 mice; *p*<0.05, K–S test). Similarly, sIPSC amplitude was also reduced in *Nrp2*^*-/-*^ mice (Fig. 4D: amplitude in pA, *Nrp2*^*+/+*^: 30.81±2.74, median=32.16, IQR=25.08-44.72, n=12; *Nrp2*^*+/-*^: 28.39±1.74, median=29.26, IQR=23.91-39.13, n=13; *Nrp2*^*-/-*^: 28.52±2.35, median=30.05, IQR=24.51-40.74, n=19; *p*<0.05, K–S test). Thus, consistent with the reduction in interneuron populations, spontaneous synaptic inhibitory inputs to CA1 pyramidal neurons is also significantly decreased in *Nrp2*^*-/-*^ mice. It is notable that while the apparent reduction in PV+ and NPY+ interneuron numbers in mice heterozygous for *Nrp2* deletion failed to reach statistical significance (Fig. 1-2), *Nrp2*^*+/-*^ mice received fewer sIPSCs than littermate *Nrp2*^*+/+*^ mice demonstrating reduced functional inhibition.

**Figure 4.**
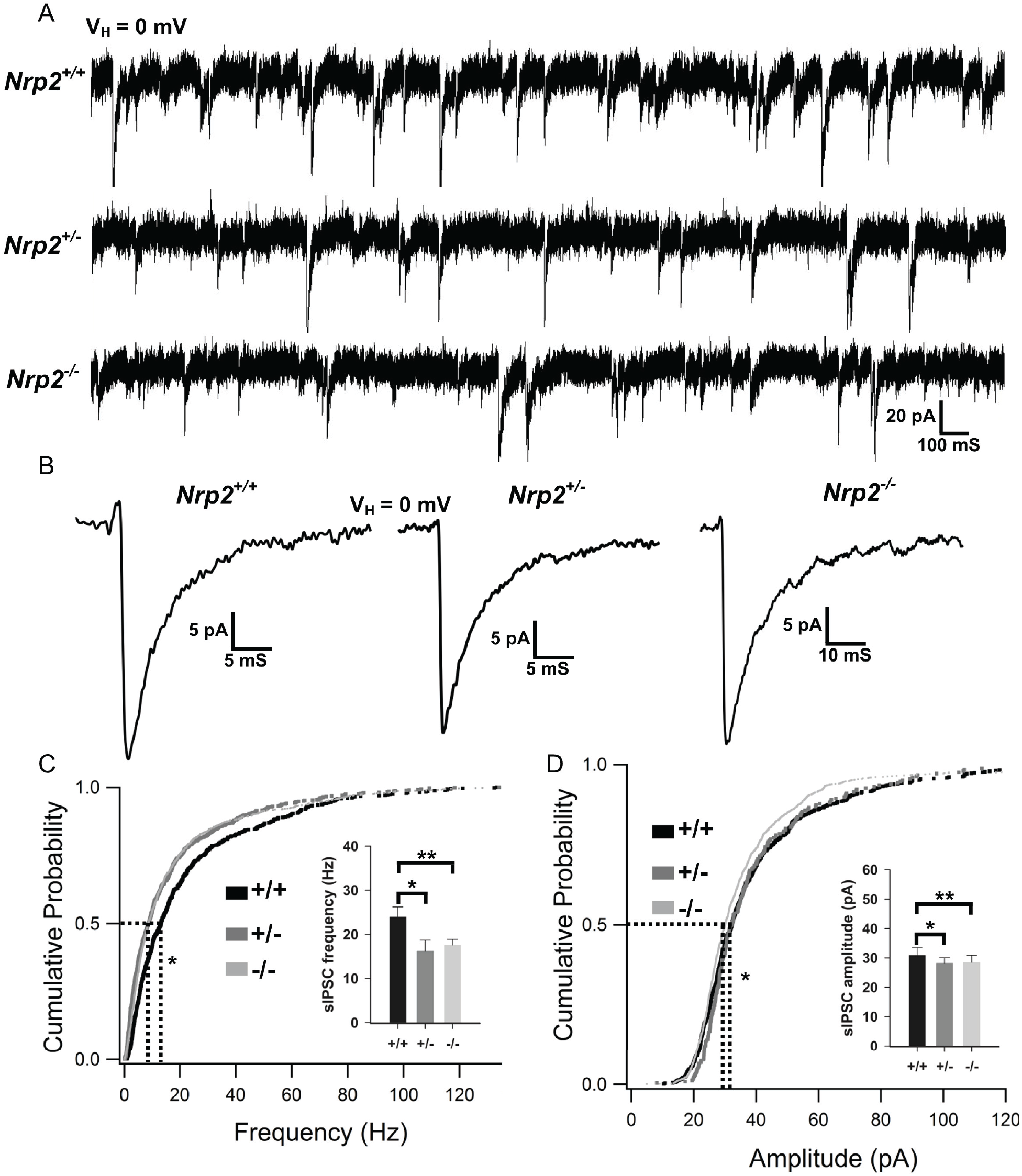
Differences in pyramidal cells sIPSC frequency and amplitude between *Nrp2* knock-outs, heterozygotes and *wild-type* littermates. A) Representative sIPSC traces from CA1 pyramidal cells of *Nrp2*^*+/+*^, Nrp2^+/-^ and *Nrp2*^*-/-*^. B) Representative average sIPSC traces from the three groups. C) Cumulative Probability plots for frequency of sIPSCs in *Nrp2*^*+/+*^ (n=12), *Nrp2*^*+/-*^ (n=13) and *Nrp2*^*-/-*^ (n=19). Inset, summary bar plot showing the frequency of sIPSCs from the three genotypes. D) Cumulative Probability plots for amplitude of sIPSCs in *Nrp2*^*+/+*^ (n=12), *Nrp2*^*+/-*^ (n=15) and *Nrp2*^*-/-*^ (n=19). * indicates p<0.05 by Kolmogorov-Smirnov test.

### Nrp2 deletion alters intrinsic physiology of CA1 Pyramidal Neurons

We next sought to determine whether deletion of Nrp2 signaling and the ensuing network reorganization impacts intrinsic active and passive properties of CA1 Pyramidal cells. Analysis of the voltage responses to graded current injections revealed a consistent reduction in frequency of action potential firing in CA1 pyramidal cells from *Nrp2*^*+/-*^ and *Nrp2*^*-/-*^ mice compared to age-matched *Nrp2*^*+/+*^ littermates (Fig. 5, n=17 cells / 4 *Nrp2*^*+/+*^ mice, 11 cells from 4 *Nrp2*^*+/-*^ mice and 13 cells from 5 *Nrp2*^*-/-*^ mice). Additionally, there was an increase in action potential amplitude in *Nrp2*^*+/-*^ and *Nrp2*^*-/-*^ relative to *Nrp2*^*+/+*^ (Fig. 5C: action potential amplitude in mV, *Nrp2*^*+/+*^: 73.50±2.00, n=16 cells/4 mice; *Nrp2*^*+/-*^: 83.23±1.64, n=10 cells/4 mice; *Nrp2*^*-/-*^: 85.82±1.58, n=11 cells/5 mice; *p*=0.001, *F*_(1,36)_=13.167, DF=36, One-way ANOVA). There was an apparent increase in spike frequency adaptation ratio in *Nrp2*^*+/-*^ and *Nrp2*^*-/-*^ mice, which failed to reach statistical significance (Fig. 5D: spike adaptation ratio, *Nrp2*^*+/+*^: 0.52±0.05, n=16; *Nrp2*^*+/-*^: 0.63±0.07, n=8; *Nrp2*^*-/-*^: 0.65±0.06 mV, n=13; *p*=0.193, One-way ANOVA). Examination of passive properties revealed that the resting membrane potential in pyramidal cells from mice lacking Nrp2 was hyperpolarized relative to *Nrp2*^*+/+*^ mice (Fig. 5E: resting membrane potential in mV, *Nrp2*^*+/+*^: -61.12±-0.82, n=17; *Nrp2*^*+/-*^: -65.40±-1.25, n=11; *Nrp2*^*-/-*^: - 64.57±-0.75, n=13; 0.004, *F*=16.388, DF=40, One-way ANOVA). However, the input resistance was not different between groups (Fig. 5F: R_in_ in MΩ, *Nrp2*^*+/+*^: 203.23±16.73, n=17; *Nrp2*^*+/-*^: 178.63±12.94, n=11; *Nrp2*^*-/-*^: 174.74±12.23, n=13; *p*=0.481, h=1.462 with 2 DF, ANOVA on Ranks). Similarly, the sag ratio, which is a measure of h-currents, was not different between groups (Fig. 5G: Sag ratio, *Nrp2*^*+/+*^: 0.84±0.01, n=12; *Nrp2*^*+/-*^: 0.84±0.01, n=10; *Nrp2*^*-/-*^: 0.84±0.01, n=13; *p*=0.927, *F*=0.0765, DF=34, One-way ANOVA). Together the reduction in firing frequency and hyperpolarized resting potential both suggest that CA1 pyramidal cell excitability is reduced in both *Nrp2*^*+/-*^ and *Nrp2*^*-/-*^ mice when compared to *Nrp2*^*+/+*^mice.

**Figure 5.**
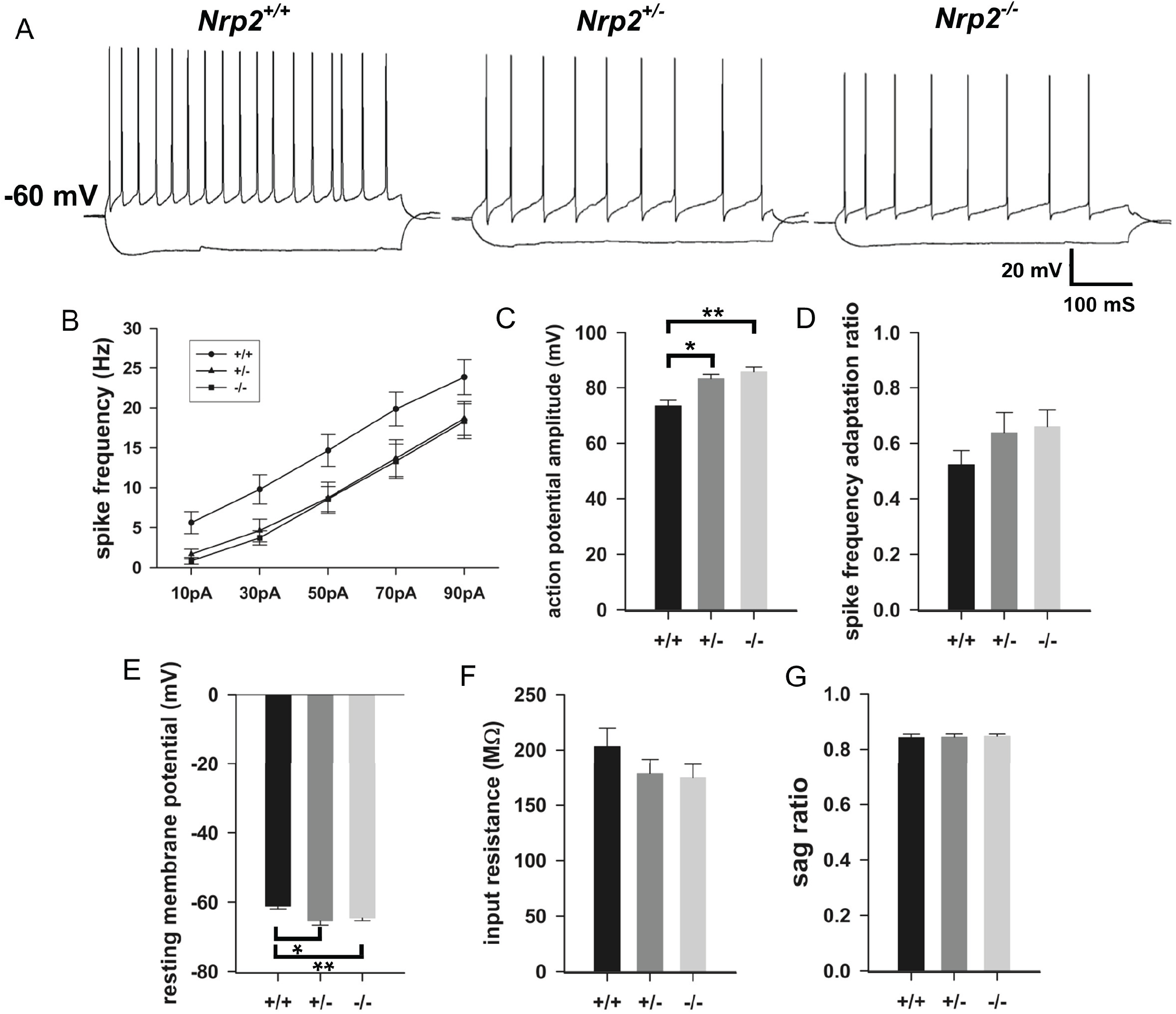
Pyramidal cells intrinsic physiology is altered by Nrp2 deletion. A) Representative current-voltage traces from *Nrp2*^*+/+*^ (left), *Nrp2*^*+/-*^ (middle) and *Nrp2*^*-/-*^ (right) CA1 pyramidal cells. B) I/F curve showing the frequencies of firing in response to 10-90 pA current injection in *Nrp2*^*+/+*^ (n=17), *Nrp2*^*+/-*^ (n=11) and *Nrp2*^*-/-*^ (n=13). Significant effect of genotype *F*_(2,38)_=4.006, *p*=0.026 by two-way repeated measures ANOVA and p<0.05 for pairwise comparison *Nrp2*^*+/+*^ and *Nrp2*^*-/-*^ using Holm-Sidak method. C-G) Summary plots of Action Potential Amplitude (C) Spike Frequency Adaptation Ratios (D), Resting Membrane Potential (E), Input Resistance (F), and Sag Ratio (G). n=16-17 for *Nrp2*^*+/+*^, 8-11 for *Nrp2*^*+/-*^ and 11-13 for *Nrp2*^*-/-*^. * indicates p<0.05 one-way ANOVA followed by pairwise comparison using Holm-Sidak method. Data presented as mean±sem.

### Nrp2 deletion accelerates development of Kainic acid (KA) induced seizures

To evaluate the impact of Nrp2 deletion and the resulting complementary changes of reduced inhibition and pyramidal cell intrinsic excitability on global hippocampal network excitability, mice were implanted with hippocampal depth electrode and injected with a single convulsive dose of KA (25mg/kg) to examine development of seizures. *Nrp2*^*-/-*^ mice exhibited a significantly shorter latency to convulsive seizures as compared to *Nrp2*^*+/+*^ mice (Fig. 6, Latency in min: *Nrp2*^*-/-*^: 4.68±1.25, median: 4.30, IQR: 3.22-5.34, n=5; *Nrp2*^*+/-*^: 16.83±3.91, median: 14.83, IQR: 10.1-18.8, n=6 and *Nrp2*^*+/+*^: 32.23±7.87, median: 30.25, IQR: 18-36.4, n=6, *p*<0.01 for *Nrp2*^*-/-*^ vs. *Nrp2*^*+/+*^, Kruskal-Wallis Rank Sum Test followed by pairwise comparison with Dunn’s test). While the seizure latency in *Nrp2*^*+/-*^ mice trended to be lower than in *Nrp2*^*+/+*^ mice the difference did not reach statistical significance. Similarly, the total duration of seizure activity measured in the first 30 minutes after KA injections was significantly longer in *Nrp2*^*-/-*^ mice as compared to *Nrp2*^*+/+*^ mice while the seizure duration in *Nrp2*^*+/-*^ mice was not different from *Nrp2*^*+/+*^ mice (Fig. 6, Time in minutes: *Nrp2*^*-/-*^: 7.19±0.73, n=4; *Nrp2*^*+/-*^: 3.08±0.98, n=5 and *Nrp2*^*+/+*^: 2.50±1.28, n=5, *p*<0.05 for *Nrp2*^*-/-*^ vs. *Nrp2*^*+/+*^; p=0.05 for *Nrp2*^*-/-*^ vs. *Nrp2*^*+/-*^, F_(2,11)_ = 5.29,, One Way ANOVA followed by pairwise comparison with Tukey’s test). Severity of behavioral seizures, scored based on the modified Racine’s scale (see methods), was also greater in *Nrp2*^*-/-*^ with four out of five mice reaching a Racine score of 5 within 10 minutes of KA injection while one out of five *Nrp2*^*+/-*^ mice and none of the *Nrp2*^*+/+*^ reached a Racine scale of 5 during the same period. The cumulative seizure score in the first 30 min post KA injection showed *Nrp2*^*-/-*^ mice had significantly higher convulsive seizures compared to *Nrp2*^*+/+*^ mice. Nevertheless, the difference between *Nrp2*^*-/-*^ mice and *Nrp2*^*+/-*^mice failed to reach statistical significance (Fig 6: *Nrp2*^*-/-*^: 23.4±1.44, median: 25, IQR: 21-26, n=5; *Nrp2*^*+/-*^: 13.3±2.17, median: 12.5, IQR: 11.2-13.8, n=6 and *Nrp2*^*+/+*^: 9.3±3, median: 7.5, IQR: 4.5-10.5, n=6, *p*<0.01 for *Nrp2*^*-/-*^ vs. *Nrp2*^*+/+*^, Kruskal-Wallis Rank Sum Test followed by pairwise comparison with Dunn’s test. Together these data reveal that *Nrp2*^*-/-*^ mice exhibit increases susceptibility and greater severity of electrographic and behavioral seizures as compared to *Nrp2*^*+/+*^ mice.

**Figure 6.**
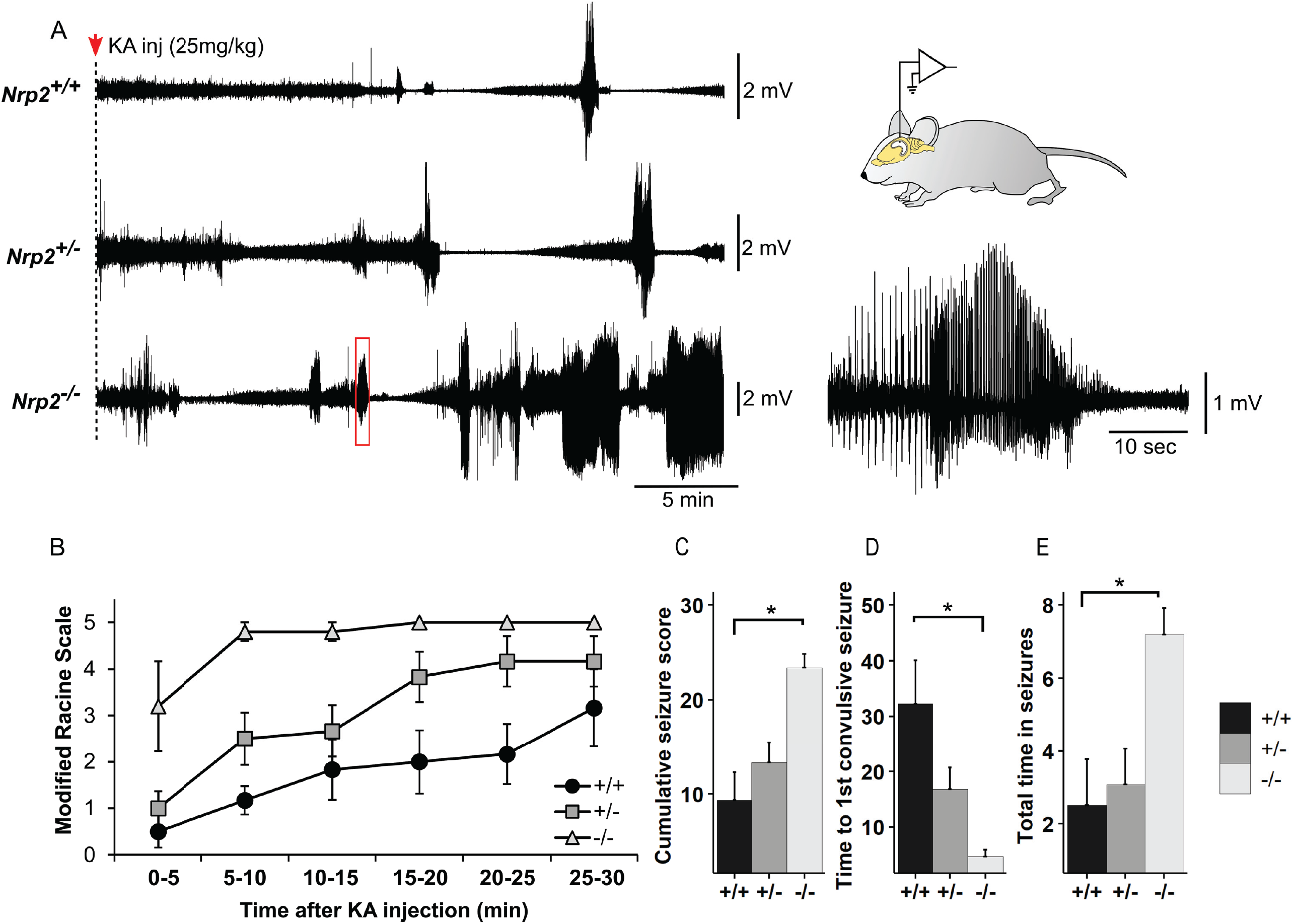
Increased susceptibility to chemically induced seizures in mice lacking Nrp2. A) Representative EEG traces show development of electrographic seizures in *Nrp2*^*+/+*^ (top), *Nrp2*^*+/-*^ (middle) and *Nrp2*^*-/-*^ (bottom) mice during the first 30 min following injection of Kainic acid (25mg/kg). Schematic of electrode placement and expanded EEG recording of seizure are illustrated on the right B) Latency to first convulsive seizure following KA inj. Note that *Nrp2*^*-/-*^ mice develop convulsive seizures significantly faster than *Nrp2*^*+/-*^, *Nrp2*^*+/+*^ mice. * indicate *p*<0.05 by pairwise comparison using Dunn test, following Kruskal-Wallis rank sum test. C. Cumulative seizures score indicates severity of behavioral seizures scored using the modified Racine scale. D) Time to first convulsive seizure following KA injection. E) Total duration of electrographic seizures in the first 30 min after KA inj shows that that *Nrp2*^*-/-*^ mice spend significantly longer duration seizing compared to *Nrp2*^*+/+*^, *Nrp2*^*+/-*^ mice. * indicate *p*<0.05 by one-way ANOVA, post-hoc Tukey’s pairwise comparison.

### Loss of Nrp2 reduces cognitive flexibility in an operant reversal-learning task

One of the hallmarks of ASD-associated behaviors is repetitive behavior ^44^, which has been suggested as result of “extreme” habit formation and inability to flexibly execute goal-directed actions ^45^. To evaluate how loss of *Nrp2* affects cognitive flexibility, we examined the performance of mice on an operant reversal-learning task (Fig.7A). All mice learned to make an operant response for food rewards; however, during the reversal task *Nrp2*^*-/-*^ and *Nrp2*^*+/-*^ mice diverged from controls in their responses following loss trials. *Nrp2*^*-/-*^ and *Nrp2*^*+/-*^ mice showed a significantly reduced lose-shift ratio compared to *Nrp2*^*+/+*^ mice (Fig. 7B; ANOVA *F*_(2,12)_= 5.25, *p*=0.0230 followed by Tukey’s test for multiple comparisons: *p*=0.0320 *Nrp2*^+/+^ vs. *Nrp2*^-/-^; *p*=0.0496 *Nrp2*^+/+^ vs. *Nrp2*^+/-^). We observed no difference between groups in responses on trials following a reward, as shown by no significant difference in the win-stay ratio (Fig. 7C; *p*>0.3 for all comparisons). Additionally, *Nrp2*-deficient mice completed a similar number of trials as control mice. These data suggest that basic appetitive motivational and reward processes were intact in *Nrp2*-deficient mice. Rather, the inability to shift responses after a loss trial from the *Nrp2*^*-/-*^ animals is indicative of behavioral inflexibility.

**Figure 7.**
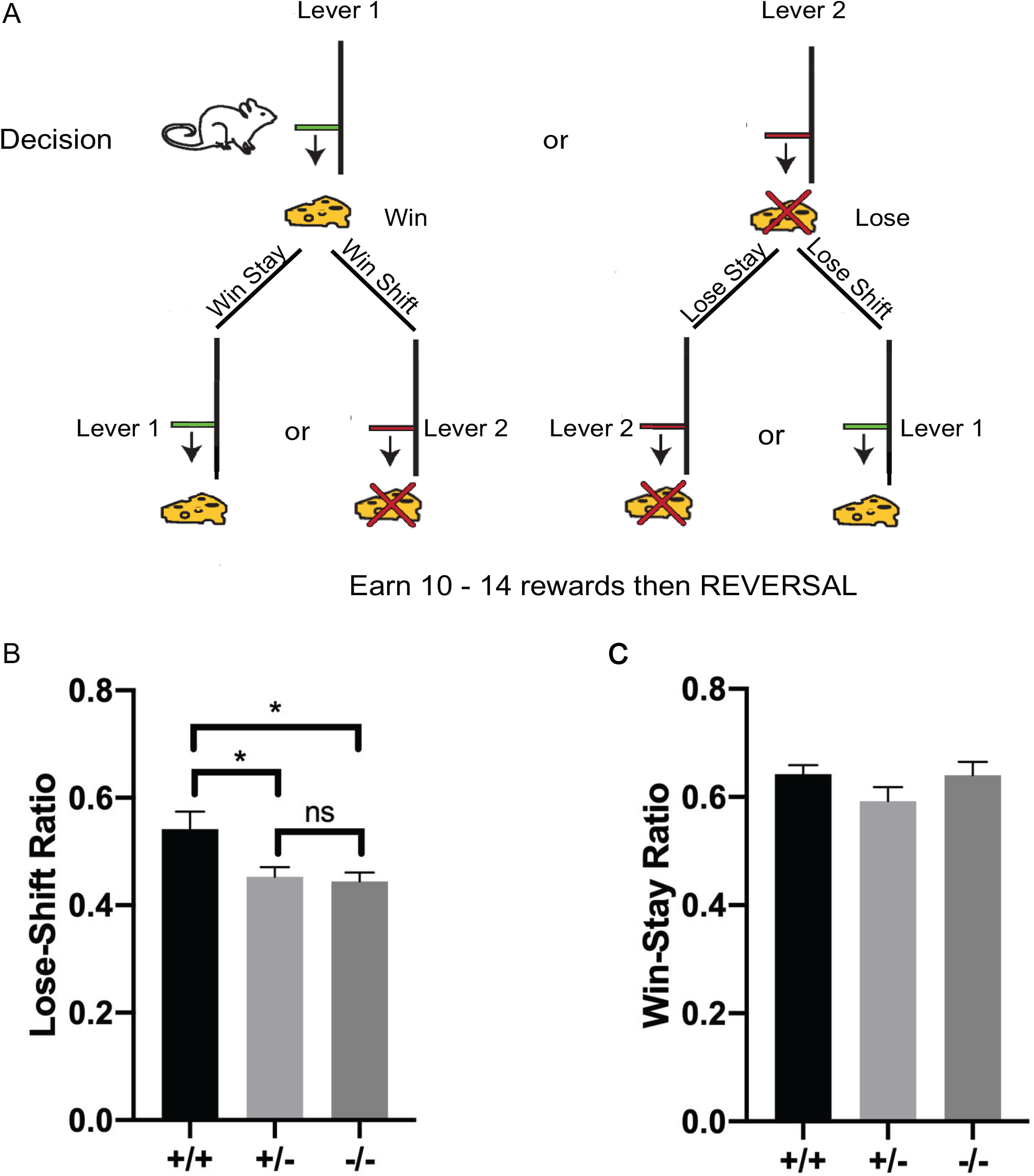
*Nrp2*-deficient mice have impaired cognitive flexibility as seen in an operant reward-based reversal-learning task. A) Schematic illustration of the reversal-learning task. Mice chose between two levers for a food reward. Responses on each trial were categorized as “win” or “loss” and “stay” or “shift” based on the previous trial’s choice and outcome. B) The lose-shift ratio was significantly reduced in *Nrp2*^*-/-*^ and *Nrp2*^*+/-*^ compared to *Nrp2*^*+/+*^ mice. C) No significant differences between genotypes were found for the win-stay ratio. Error bars are ± SEM; one-way ANOVA, post-hoc Tukey: \**p*=0.0320 +/+ vs. -/-, \**p*=0.0496 +/+ vs. +/-, and ns *p*=0.9669, n=5 animals/genotype.

## Discussion

It is well established that alterations in GABAergic circuits impact a wide variety of neurobiological functions leading to neurological disorders, including the highly comorbid conditions of ASD and epileptic seizures ^1,17,46^. However, in the contribution of perturbations in different classes of GABAergic interneurons to the pathophysiological mechanisms and the associated behavioral phenotypes observed in autism and epilepsy is not well understood. In our study, we investigated a loss-of-function genetic mouse model, specifically targeted to the *Nrp2* locus, which has known mutations linked to autistic patients ^22,23^, to determine the cellular, physiological and behavioral aspects of Nrp2’s contribution to epileptic seizures and ASD-like phenotypes when single versus double copies of *Nrp2* is deleted. We demonstrated that Nrp2 is required for appropriate maturation of synaptic inhibition and regulates the density of of multiple hippocampal interneuron types; largely reducing SOM+ cell numbers in all CA subregions and DG of the hippocampus. We also showed that both Nrp2 heterozygous and homozygous mutants have reduced intrinsic excitability of CA1 pyramidal neurons, which could indicate an innate compensatory mechanism to restrain excessive excitation in the presence of reduced synaptic inhibition and interneuron numbers. We identified that the loss of diverse interneuron classes, which support inhibition across multiple dendritic zones ^42^, was associated with increase susceptibility to kainic acid induced seizures in the *Nrp2* mutant animals as compared to littermate controls. Lastly, we challenged the *Nrp2* mutant mice in a reversal-learning task, which is used to measure cognitive flexibility ^47^. The ability to make a flexible choice is often compromised in ASD individuals, who are frequently dominated by restrictive and repetitive behaviors ^48^. There is increasing evidence that intact hippocampus circuit function is necessary to respond in an appropriately flexible manner, and disruptions render behavior habitual and inflexible ^49,50^. Interestingly, both *Nrp2*^*+/-*^ and *Nrp2*^*-/-*^ showed impaired cognitive flexibility as illustrated with significantly lower lose-shift ratios compared to *Nrp2*^*+/+*^ control animals. Taken together, our findings provide novel insights into the cellular and physiological basis contributing to the behavioral phenotypes observed in heterozygous and homozygous *Nrp2* mutant mice, and may serve as a useful model for studying the molecular mechanisms underlying developmental processes contributing the propensity for ASD and epilepsy comorbidity observed clinically.

### Impact of Nrp2 deletion on hippocampal inhibition and seizure susceptibility

Our finding that global *Nrp2* deletion causes significant decreases in three distinct interneuronal classes, PV+, SOM+ and NPY+ neurons in the hippocampal CA1 supports the role for Nrp2 in interneuron development ^14,41^. In contrast to prior reports ^19^, we find that SOM+ neurons show most extensive reduction in all major hippocampal subregions while PV+ and NPY+ showed selective decrease in CA1. It is likely that the reduction in hippocampal interneuronal population following Nrp2 deletion arises from disruption of migration of interneuron precursors from the MGE during development ^41^. However, the decrease in interneuron population identified here differs from the limited cell-specific depletion of NPY+ neurons in *Nrp2*^*-/-*^ mice or lack of interneuron reduction in Sema3F knockout animals reported previously ^19,25^. It is possible that methodological differences, particularly, age of the animals examined may underlie these differences in observations as migration of progenitors may still be an active process in younger animals..

In keeping with the extensive interneuron loss, we find a significant reduction in the frequency of inhibitory synaptic inputs to CA1 pyramidal neurons in mice lacking Nrp2. Unlike the cell loss which was observed only in *Nrp2*^*-/-*^ mice, the functional reduction in inhibition was observed in *Nrp2*^*+/-*^ littermates suggesting that, in addition to decrease in interneuron populations, circuit level plasticity in Nrp2 deficient mice may also impair inhibition. In this regard, Nrp2 deletion is known to increase dendritic spines and excitatory inputs to glutamatergic neurons and regulate synapse formation and plasticity ^12,30^. Our sIPSC recordings, conducted in glutamate receptor antagonists to limit the contribution of known changes in the excitatory inputs ^30^, identify that reducing Nrp2 signaling impairs basal inhibition. Interestingly, CA1 pyramidal neurons in Nrp2 deficient mice rest at more hyperpolarized potentials and have reduced firing in response to depolarizing current injections indicating that the changes in excitation inhibition (E/I) balance brought about by the increased excitatory drive ^30,34^ and reduce inhibition (Fig. 4) may lead to compensatory metaplastic changes in intrinsic excitability. The reduction in resting membrane potential and excitability were not accompanied by changes sag ratio, a measure of h-currents, or input resistance and the channel mechanisms underlying these changes need further investigation. Despite the potential compensatory decrease in cell intrinsic excitability, we find that the E/I imbalance reduces the latency to chemically induce seizures and increases seizure severity demonstrating increased epileptogenicity in *Nrp2*^*-/-*^ mice. Although there were trends suggesting increased seizure susceptibility in *Nrp2*^*+/-*^ mice, these did not reach statistical significance. The relative maintenance of interneuron numbers and seizure susceptibility in *Nrp2*^*+/-*^ mice, despite impairments in intrinsic excitability, synaptic inhibition and cognitive flexibility indicate some network-level compensation which, while sufficient to prevent epilepsy, may compromise finer cognitive processing.

### Hippocampal-dependent cognitive flexibility in ASD phenotypic analysis

We assessed the ability of *Nrp2*^*+/+*^, *Nrp2*^*+/-*^ and *Nrp2*^*-/-*^ mice to perform an operant reward-based deterministic reversal-learning task. In this task, mice use reinforcement history to guide their choice of actions. The optimal strategy is to choose the opposite lever following a non-rewarded “loss” trial and to repeat the same action after a rewarded “win” trial. We found that both *Nrp2*^*+/-*^ and *Nrp2*^*-/-*^ mice made fewer shifts to the opposite lever following a loss trial, as shown by a reduced lose-shift ratio compared to control mice. Rather, nrp2-deficient mice continued to respond on the non-rewarded lever, which suggests an impairment in cognitive flexibility. Specifically, the repetition of non-rewarded responses following loss trials is a form of response perseveration, which is a well-known trait of autism ^45^. In a previous study ^20^, we observed a similar form of response perseveration in *Nrp2* null mice, in which mice displayed enhanced self-directed grooming. In contrast to responses following loss trials, Nrp2-deficient mice did not differ from controls in their responses following rewarded “win” trials, suggesting that basic appetitive motivational processes are not disturbed.

Cognitive inflexibility in reversal learning tasks have also been shown in rats, Rhesus monkeys and humans following lesions of the hippocampus ^49,51-53^. In one study, faster reversal learning rates were shown to be associated with an increased ratio of PV+ interneurons to granule cells in the hippocampus ^54^. Consistent with these findings we found fewer PV+ as well as other interneuron subtypes in the hippocampus of *Nrp2*^*-/-*^ mice compared to both *Nrp2*^*+/+*^ and *Nrp2*^*+/-*^ mice. *Nrp2*^*+/-*^ and *Nrp2*^*-/-*^ mice both showed deficits in the reversal-learning behavior task. While *Nrp2*^*+/-*^ mice did not differ in the number of PV+ hippocampal interneurons, they did have significantly fewer SOM+ interneurons in the entire hippocampus compared to *Nrp2*^*+/+*^ mice. Future study is required to determine the roles of specific interneuron subtypes in the hippocampus and their relation to reversal-learning.

Both inhibitory interneuron deficiency in the hippocampus and behavioral inflexibility have been associated with ASD ^45,55-57^. “Inflexible adherence to routines” is included as part of the DSM V requirements for diagnosing autism ^44^. In a reversal-learning task, individuals with autism had difficulty maintaining reversals compared to controls ^57^. Similarly, mice with a mutation an autism candidate gene, NrCam, experienced deficiencies in reversal-learning but not acquisition learning in a Morris water maze test ^58^. Here, we have demonstrated that mice with deficiencies in the autism candidate gene *Nrp2* have deficits in reversal learning strategies leading to impaired cognitive flexibility.

In summary, our studies identify that deletion of Nrp2, a gene with known polymorphisms in patient populations with autistic syndromes, contributes to a broad reduction in PV+, SOM+ and NPY+ interneuron numbers in the hippocampal CA1 and compromises synaptic inhibition. We demonstrate that these circuit level disruptions in inhibition contribute to increased epileptogenicity and impaired cognitive flexibility, a hallmark of autism spectrum disorders. Our findings in mice with global Nrp2 deletion provide a strong foundation for future studies examining how interneuron specific and developmental deletion of Nrp2 can impact cortical and hippocampal inhibitory circuit formation and their role in neuropsychiatric disorders. Taken together, our results indicate that reduced Nrp2 signaling impacts development of interneurons circuits contributing to autism-epilepsy comorbidity.

## Acknowledgements

The authors would like to thank Emma D. Eisenberg for her help in illustrating the schematic diagrams of the hippocampus in the manuscript. This work is supported by the NJ Governor’s Council for Medical Research and Treatment of Autism: CAUT17BSP011 to V.S. and T.S.T., CAUT17BSP022 to T.S.T. and M.W.S., Rutgers Brain Health Initiative Pilot Grants to T.S.T. and V.S., NIH R01 NS069861 and NS097750 to V.S., and NSF/IOS: 1556968 to T.S.T.

## Supplemental Figure Legend

**Figure S1**. Schematic drawings indicating subregions of the mouse hippocampus used for the interneuron cell counts in the *Nrp2KO*^+/+^, *Nrp2KO*^+/-^ and *Nrp2KO*^-/-^ mice. Ten sections were obtained at 200 µm intervals through the entire anterior-posterior hippocampus. Representative images shown include the anterior (A), middle (B), and posterior (C) sections of the hippocampus. All cells whose full nuclei are within these regions were counted and averaged over the 10 sections.

