## Supplementary figures and images for "Reduced Hippocampal Inhibition and Enhanced Autism-Epilepsy Comorbidity in Mice Lacking Neuropilin 2"

### Supplemental Figure S1

Supplemental Fig. S1

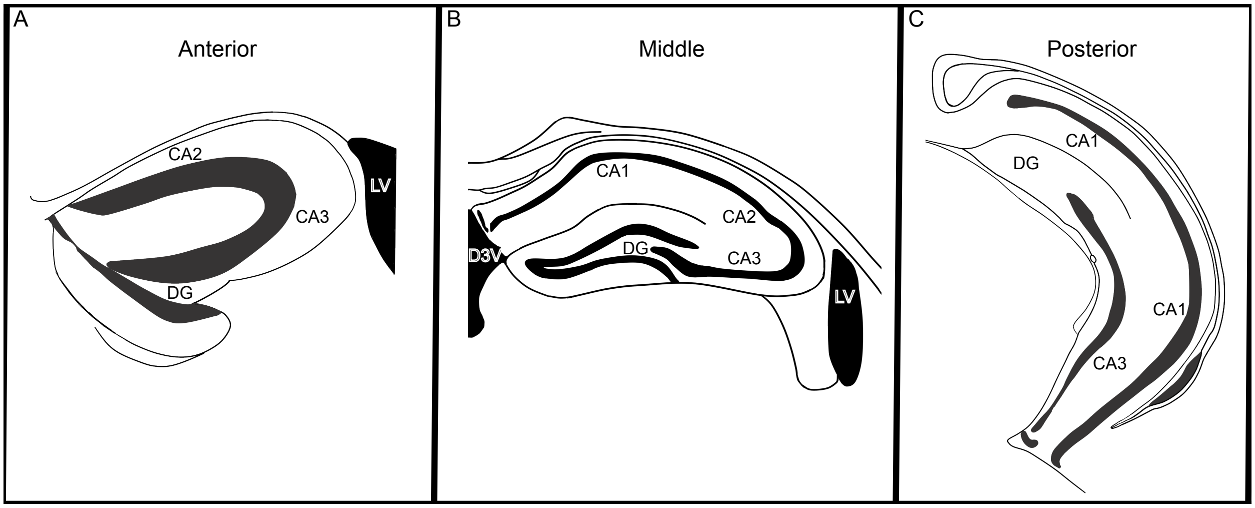
